# *Leishmania mexicana* Promotes Pain-reducing Metabolomic Reprogramming In Cutaneous Lesions

**DOI:** 10.1101/2022.08.09.503319

**Authors:** Greta Volpedo, Timur Oljuskin, Blake Cox, Yulian Mercado, Candice Askwith, Nazli Azodi, Sreenivas Gannavaram, Hira L. Nakhasi, Abhay R. Satoskar

**Affiliations:** Department of Microbiology, The Ohio State University, Columbus, OH 43210, USA; Department of Pathology, Wexner Medical Center, The Ohio State University, Columbus, OH 43210, USA; Animal Parasitic Disease Lab, Agricultural Research Service, USDA, Beltsville MD, USA; Department of Neuroscience, The Ohio State University, Columbus OH 43210, USA; Division of Emerging and Transfusion Transmitted Diseases, CBER, FDA, Silver Spring, MD, USA.

**Keywords:** Analgesia, cutaneous leishmaniasis, metabolic reprogramming, metabolomics

## Abstract

Cutaneous leishmaniasis (CL) is characterized by extensive skin lesions associated with an aggressive inflammatory reaction. Despite the extensive inflammation, CL lesions are usually painless, indicating that *Leishmania* infection may trigger anti-nociceptive activities in the infected tissues. To this date, the molecular mechanisms responsible for this clinical phenomenon have not been identified. Through an untargeted metabolomic analysis by mass spectrometry, we found enriched anti-nociceptive metabolic pathways in mice infected with *Leishmania* (*L.*) *mexicana.* In particular, endogenous purines were elevated at the lesion site during chronic infection, as well as *in vitro* in infected macrophages, compared to non-infected mice. These purines have known anti-inflammatory and analgesic properties by acting through adenosine receptors and inhibiting transient receptor potential channels of the vanilloid subtype 1 (TRPV1). Additionally, purine metabolites can promote interleukin (IL)-10 production, with a subsequent decrease in inflammation and pain sensitivity. We also found arachidonic acid metabolism enriched in the ear lesions compared to the non-infected controls. Arachidonic acid is a metabolite of anandamide (AEA) and 2-arachidonoylglycerol (2-AG). These endocannabinoids act on cannabinoid receptors 1 and 2 and TRPV1 channels to exert anti-inflammatory and analgesic effects. Our study provides the first evidence of metabolic pathways upregulated during *L. mexicana* infection that may mediate anti-nociceptive effects experienced by CL patients and identifies macrophages as a source of these metabolites.

**Graphical abstract:** *L. mexicana* infection promotes the production of purines, as well as endocannabinoid mediators, which could act on different channels of dorsal root ganglia neuron to inhibit nociception.

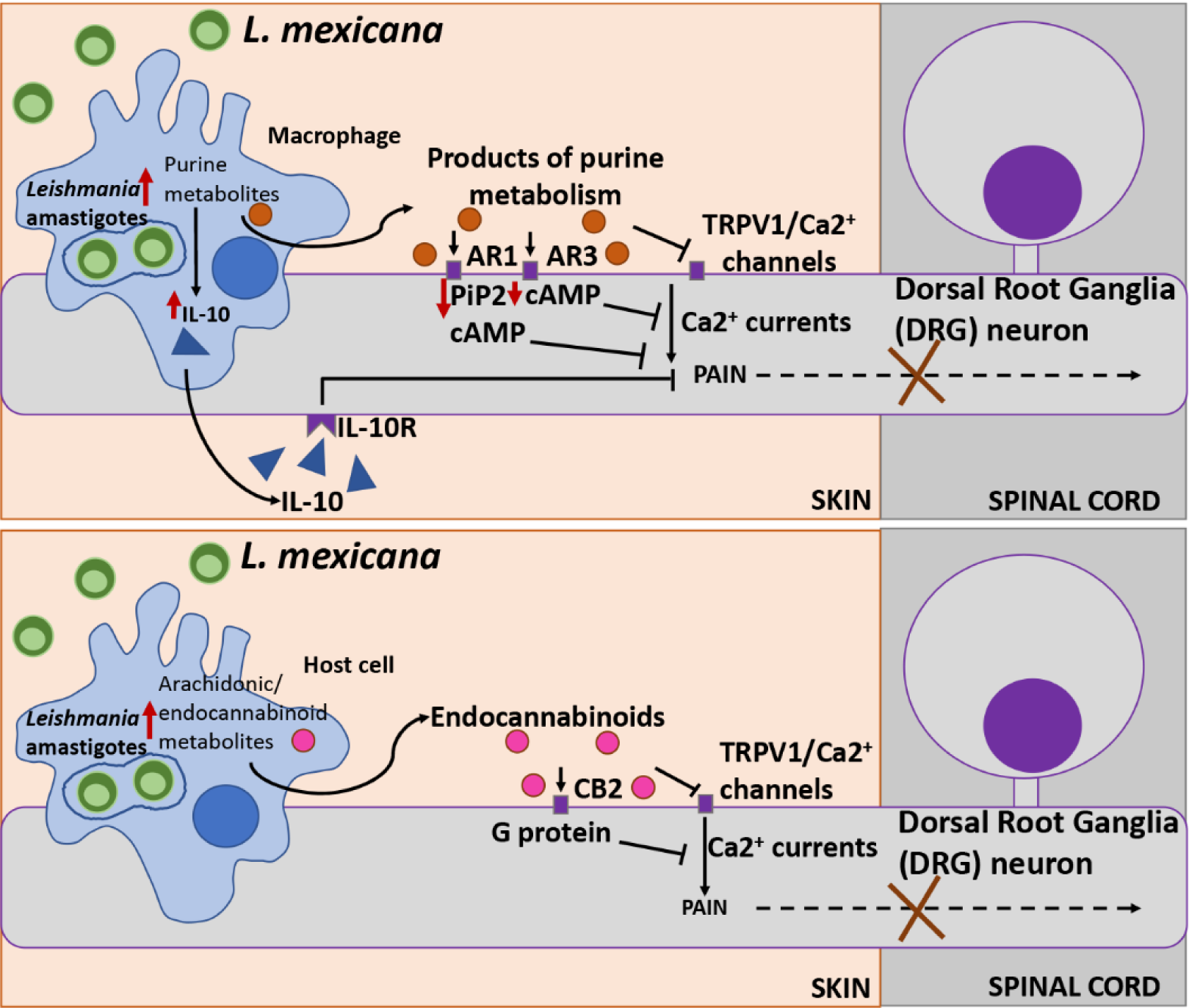

## INTRODUCTION

Cutaneous leishmaniasis (CL) is caused by the protozoan parasite *Leishmania* and it is the most common form of the disease with an incidence of 1 million new cases every year^1, 2^. Depending on the causative species, the clinical manifestations of CL can range from self-healing skin lesions to diffuse ulcers that can persist for months and require treatment^3^. *L. mexicana* is a New World CL strain pervasive in Central and South America, which can cause chronic non-healing skin lesions on different parts of the body^4, 5^. Interestingly, CL lesions, including those caused by *L. mexicana*, are usually painless in humans^6–8^, a clinical phenomenon that has not received much attention from the scientific community. This is an intriguing phenotype as inflammatory lesions, such as lesions caused by *Staphylococcus aureus*^9^, *Herpes* virus^10^, and *Varicella zoster* virus^11^, are generally painful. CL lesions can range between 0.5 and 3 cm in diameter^12^ and present with chronic inflammation and infiltration of lymphocytes, plasma cells, and macrophages^13, 14^. Despite the extensive inflammation, CL lesions are usually painless, indicating that *Leishmania* infection may trigger anti-nociceptive activities in the infected tissues. Interestingly, pain is restored in human CL lesions after secondary bacterial infections, which are common following ulceration^12, 15^. This suggests that any mediators preventing pain produced in the primary lesions are overridden by other signals following a secondary infection and that nociception at the lesion site might be due to a pathogen-specific mechanism.

Interestingly, CL infections seem to be painless from the sand fly blood meal to the development and progression of the lesions^6–8^, which could indicate an intricate host-parasite interaction that may favor the establishment and spread of infection. Analgesia could delay the infected individual from seeking medical attention, allowing for the infection to establish and spread to other areas of the body and to other individuals via the sand fly vector. This suggests that there might be a broader biological advantage in having painless invasion and painless lesions. For these reasons, it is imperative to continue exploring this understudied phenomenon to elucidate the molecular mechanisms causing lack of pain at the lesion site.

Several studies have been conducted in mice infected with *L. major*, an Old World cutaneous strain, to explore the role of cytokines in CL-mediated analgesia^16–21^, as it is well established that these mediators can modulate pain at the neural level. Nociceptive neurons express cytokine receptors, which can modulate neuronal excitability and pain signaling^22^. For instance, interleukin (IL)-10, IL-13, and IL-4 are endogenous cytokines produced in great quantities in *Leishmania* susceptible models and are known to have analgesic properties^6^. On the other hand, pro-inflammatory cytokines have been shown to cause pain by increasing excitability of nociceptive neurons^23^. However, studies focusing on the role of cytokines in *L. major*-mediated anti-nociception have shown rather contradicting results in pre-clinical models^16–21^.

Even when analgesia was observed, changes in the expression of cytokines alone did not seem to be responsible for this phenotype^16^. These observations led us to hypothesize that other non-immunological mediators produced or upregulated during *Leishmania* infection may directly mediate anti-nociception at the lesion site. For instance, different metabolites have been previously explored for their analgesic roles. In particular, endocannabinoids^24^, protectins^25, 26^, maresins^27, 28^, resolvins^28^, and lipoxins^29^, among others, have been shown to have anti-nociceptive properties by directly acting on nociceptor neurons.

In order to acquire a complete and unbiased understanding of the mediators produced during infection, we used an untargeted metabolomics approach to identify novel *Leishmania*-induced mediators with analgesic properties. Furthermore, we investigated pain responses during *L. mexicana* infection, prevalent in geographically distinct endemic areas and presenting different disease immunology and pathology compared to *L. major*^30–33^. In particular, *L. mexicana* lesions are not self-resolving^4, 5^, contrary to *L. major* lesions, and would require a prolonged analgesic action.

In this study, we show for the first time that mice infected with *L. mexicana* display enriched metabolites with anti-nociceptive properties at the infected tissue. In particular, endogenous purines, arachidonic acid, and endocannabinoid metabolites were upregulated in the lesions of *L. mexicana*-infected mice compared to their uninfected controls and could lead to the lack of pain observed in CL patients. Exploring the metabolic-neuro-immune interaction in CL, with a particular focus on macrophages and nociceptive neurons, could elucidate the anti-nociceptive mechanism/s leading to analgesia in CL patients.

## RESULTS

### *L. mexicana* infection results in metabolic differences compared to naïve skin tissue

In order to gain a comprehensive understanding of the mediators produced during infection, as well as those that might play a role in *L. mexicana*-mediated anti-nociception, we performed untargeted mass spectrometry on the ear lesions of C57BL/6 mice chronically infected intradermally with *L. mexicana* (Fig. 1a). Our goal was to screen for enrichment of mediators with known anti-nociceptive properties in infected, compared to non-infected ear tissue. Figure 1b shows representative images of the ears of naïve and *L. mexicana*-infected mice at harvest. As expected, infected mice showed a large lesion at 12 weeks post infection, indicated by a black arrow (Fig. 1b).

**Figure 1.**
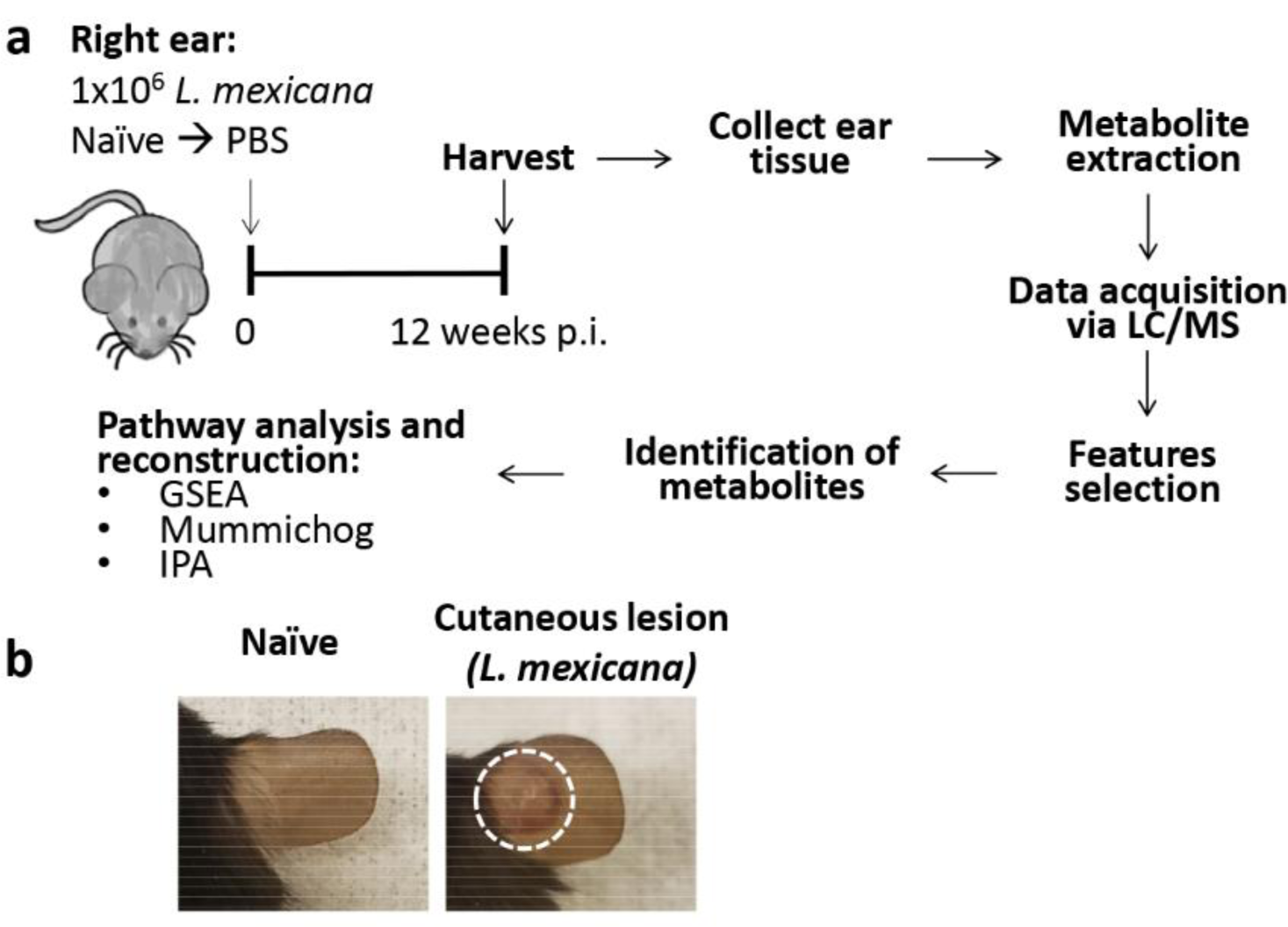
Experimental design for metabolomic analysis to detect mediators of analgesic activity following *L. mexicana* infection. **a)** Experimental design of infection, sample collection, and analysis. **b)** Representative images of the ears of naïve and *L. mexicana*-infected mice at 12 weeks post infection. Abbreviations: LC/MS: liquid chromatography / mass spectrometry; GSEA: gene set enrichment analysis; IPA: ingenuity pathway analysis.

Using volcano plots (Fig. 2a), we selected significant features from the positive and negative modes of mass spectrometry analysis. The peaks identified with the positive mode correspond to protonated molecules, while those identified with the negative mode correspond to deprotonated analytes. A fold change threshold (x) 2 and t-tests threshold (y) 0.05 were set for the volcano plots (Fig. 2a). We have also used partial least squares-discriminant analysis (PLS-DA), a supervised multivariate analytic approach, to identify unique molecular pathways enriched during *L. mexicana* infection (Fig. 2b). Not surprisingly, our analysis revealed distinct metabolic signatures in the ear tissue of naïve, compared to infected mice, highlighting a significant *Leishmania*-specific metabolic shift.

**Figure 2.**
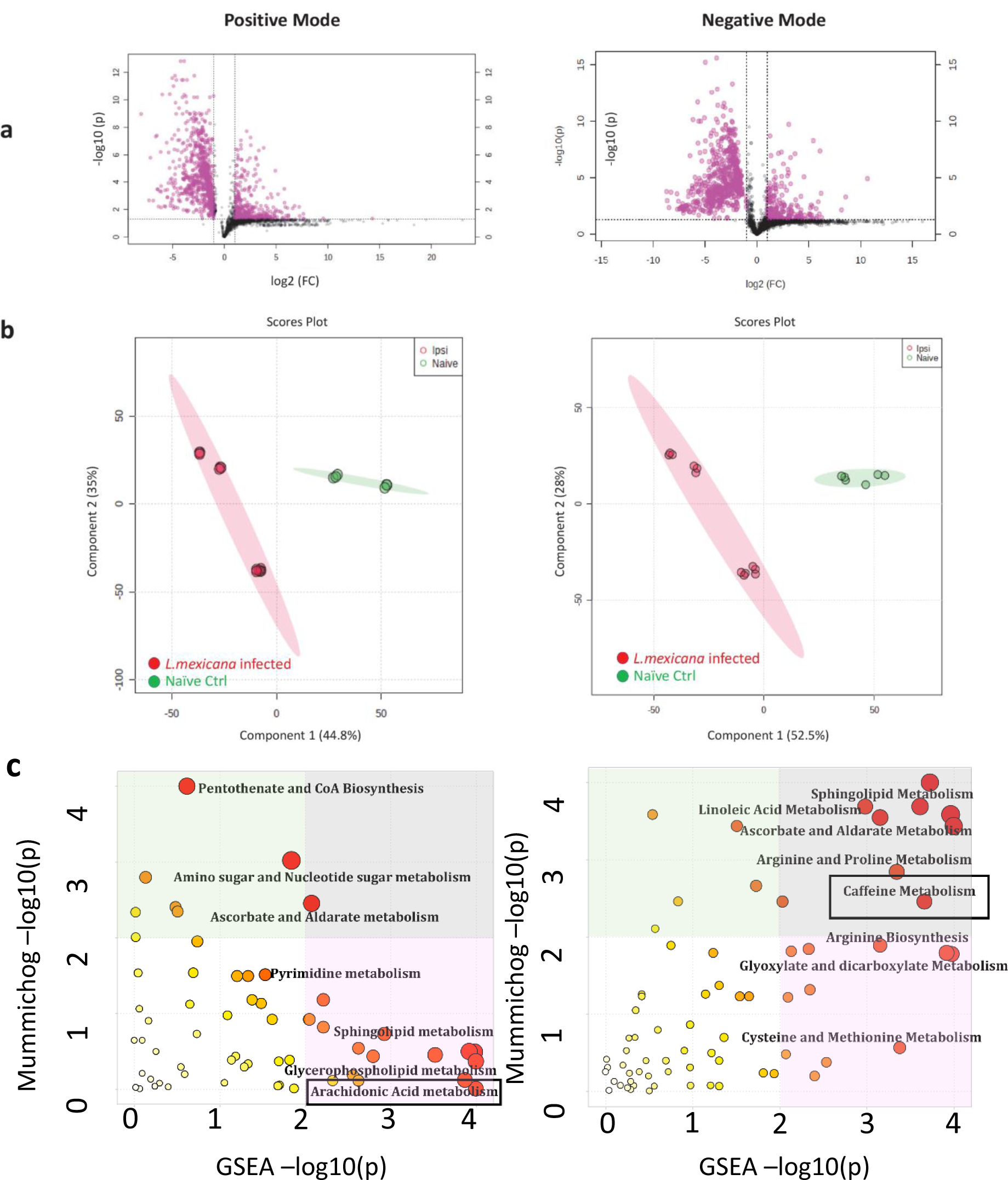
Infection with *L. mexicana* leads to enriched caffeine and arachidonic acid metabolism in cutaneous lesions. Normalized data from ear tissue of C57BL/6 mice infected with *L. mexicana* (12 weeks post infection) and naïve controls was used to perform statistical analysis. **a)** Features selected by volcano plot from positive and negative modes of LC/MS with log-transformed fold change threshold (2-fold, x-axis) and t-tests threshold (0.05, y-axis). **b)** Score plots between the selected PLS-DA from positive and negative modes are shown. **c)** The Integrated MS Peaks to Paths plot summarizes the results of the Fisher’s method for combining mummichog (y) and GSEA (x) p-values from the positive and negative mode data sets, indicating the metabolic pathways enriched. The size and color of the circles correspond to their log-transformed combined p-values. Large and red circles are considered the most perturbed pathways. The colored areas show the significant pathways based on either GSEA (pink) or mummichog (turquoise), and the purple area highlights significant pathways identified by both algorithms. Highlighted by a black box are arachidonic acid metabolism for the positive mode, and caffeine metabolism for the negative mode.

### *L. mexicana* infection leads to enrichment of purine metabolism *in vivo*

In order to identify specific anti-nociceptive metabolic pathways enriched in infected mice, we used both mummichog and gene set enrichment analysis (GSEA) analysis.

Mummichog analysis uses an algorithm that predicts functional activity bypassing metabolite identification, while GSEA requires the metabolites to be identified before pathway/network analysis. We used the Integrated MS Peaks to Pathways plot to summarize the results of the Fisher’s method for combining mummichog (y) and GSEA (x) p-values from the positive and negative mode data sets, indicating several enriched pathways at the lesion site (Fig. 2c). In particular, the negative mode data set showed enriched caffeine metabolism in infected vs. uninfected mice (Fig. 2c, black box). Based on the t scores for the matched metabolites, caffeine metabolism was upregulated in infected mice. This finding was surprising as no caffeine was administered in this study. Further investigating this pathway, we found that several endogenous purines, which are analogous to metabolites produced by the caffeine metabolism, were highly upregulated during infection. In particular, xanthosine and 5-Hydroxyisourate were significantly up-regulated in the infected group, as identified using Medscape (Figure 3). Caffeine, purines, and their metabolites have been previously implicated in anti-nociception^34, 35^ and could therefore provide a mechanism for the lack of pain reported by CL patients.

**Figure 3.**
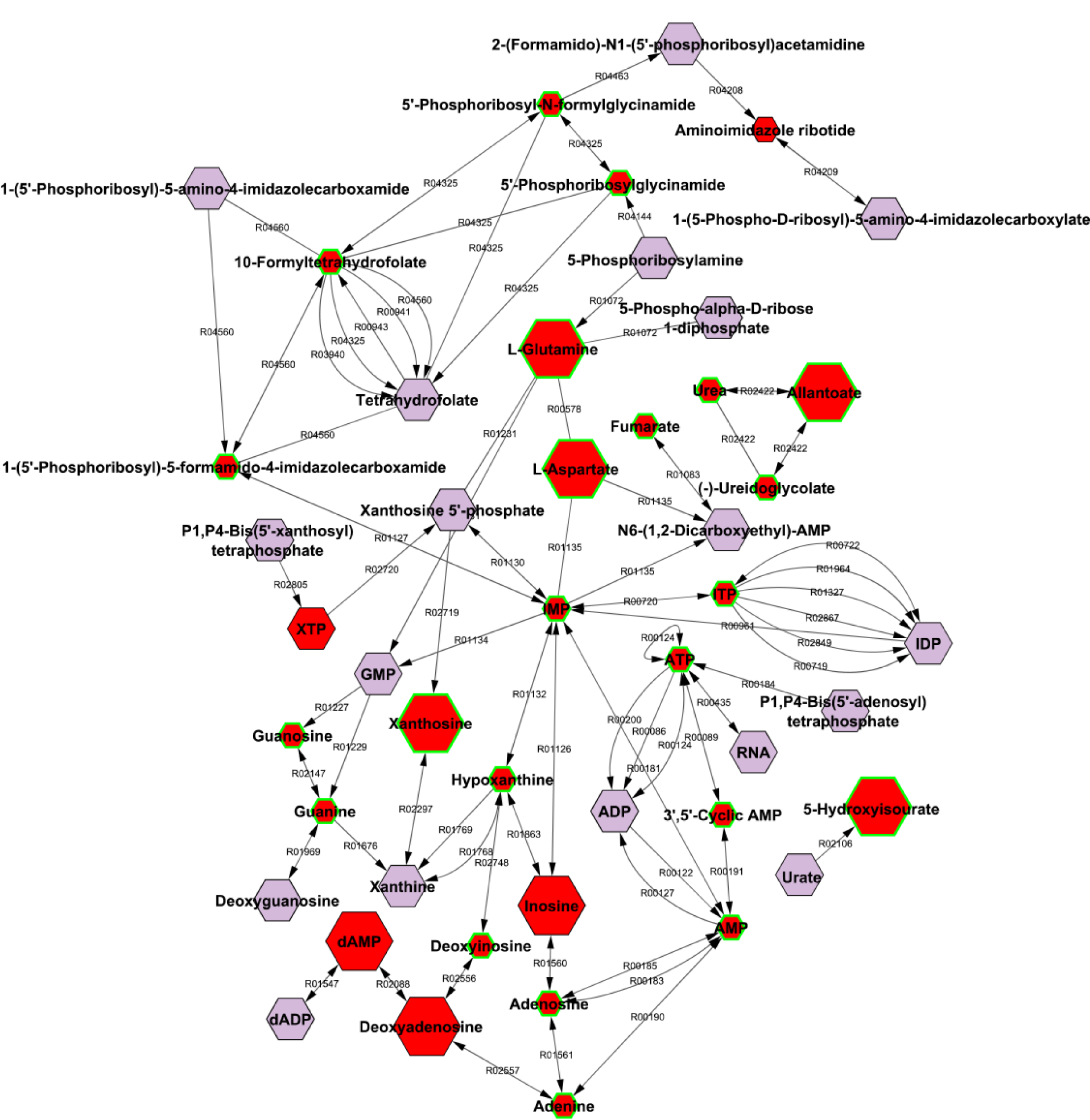
Infection with *L. mexicana* leads to enriched purine metabolism. Normalized data from ear tissue of C57BL/6 mice infected with *L. mexicana* (12 weeks post infection) and naïve controls was used to perform integrative compound-compound network analysis with Metscape. Endogenous purines analogous to those produced in caffeine metabolism are included in this graph. Larger hexagons represent up-regulation, while smaller hexagons represent down-regulation. Red hexagons represent compounds detected in the data set, while hexagons with a green outline represent statistically significant metabolites.

### *L. mexicana* infection leads to enrichment of arachidonic acid and endocannabinoid metabolites

Ear tissue of infected mice analyzed in the positive mode data set revealed additional metabolic pathways with analgesic properties, which were activated during infection. Based on GSEA analysis, a pathway of interest that was enriched in the infected ears of chronically infected mice was arachidonic acid metabolism (Fig. 2c). Based on the t scores for the matched metabolites, arachidonic acid metabolism was activated in infected mice. Arachidonic acid metabolism is closely related to endocannabinoid metabolism, known to exert potent analgesic functions^24^. In particular, we observed increased levels of diacylglycerol and monoacylglycerol, precursors of arachidonic acid, as well as phosphatidylethanolamine and phosphatidylcholine, both precursors of the endocannabinoid anandamide (AEA), as well as phosphatidylinositol-3-phosphate, a precursor of the endocannabinoid 2-arachidonoylglycerol (2-AG) (Table 1).

**Table 1.**
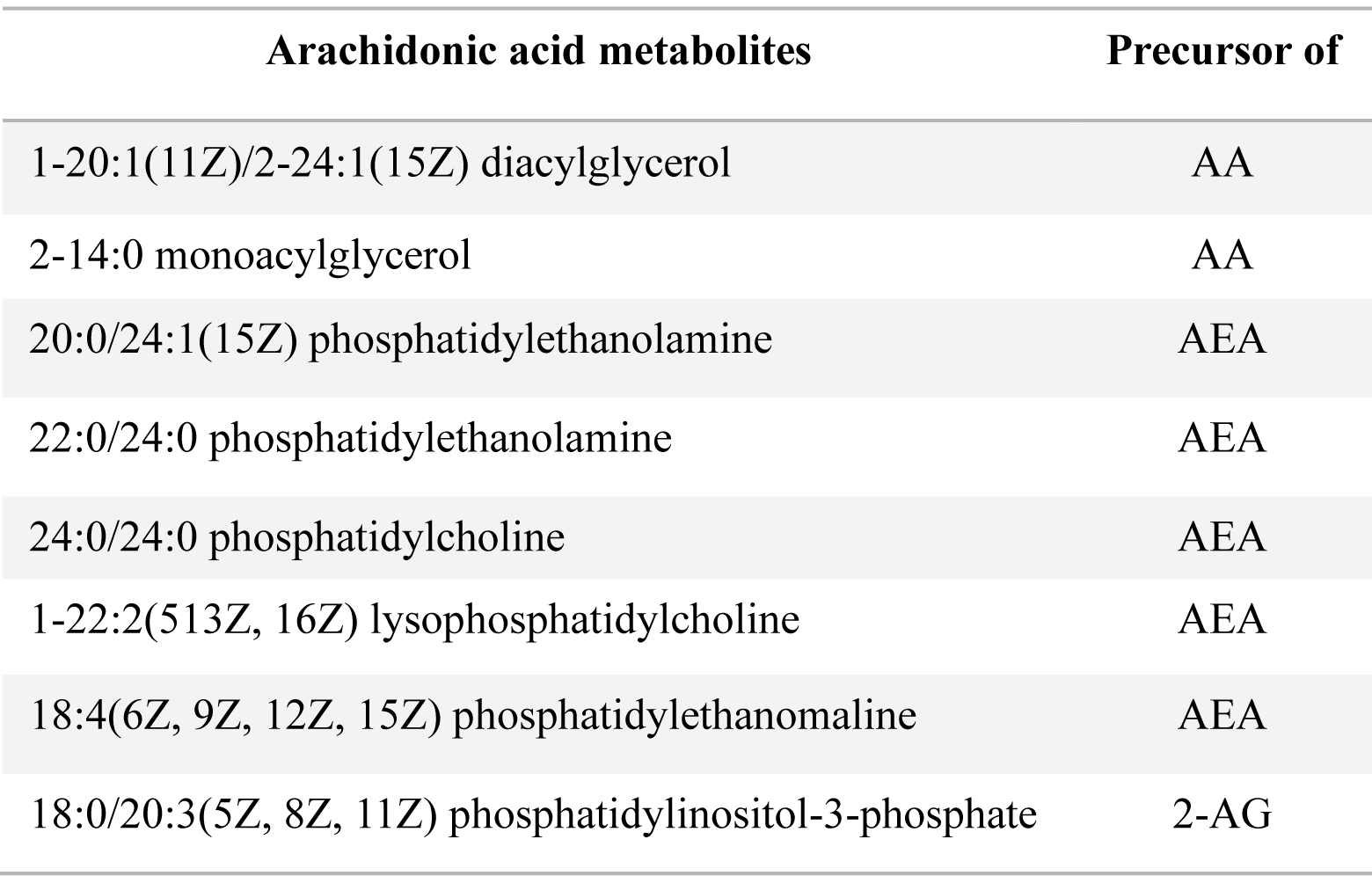
Significantly upregulated arachidonic acid metabolites in ear tissue of mice infected with L. mexicana, compared naïve controls. Normalized data from ear tissue of C57BL/6 mice infected with L. mexicana (12 weeks post infection) and naïve control was used to perform pathway analysis. Mummichog and GSEA analyses showed arachidonic acid metabolites were significantly elevated in the cutaneous lesion of L. mexicana-infected mice, compared to naïve controls. Abbreviations: AA: arachidonic acid; AEA: anandamide; 2-AG: 2-arachidonoylglycerol.

| Arachidonic acid metabolites | Precursor of |
| --- | --- |
| 1-20:1(11Z)/2-24:1(15Z) diacylglycerol | AA |
| 2-14:0 monoacylglycerol | AA |
| 20:0/24:1(15Z) phosphatidylethanolamine | AEA |
| 22:0/24:0 phosphatidylethanolamine | AEA |
| 24:0/24:0 phosphatidylcholine | AEA |
| 1-22:2(513Z, 16Z) lysophosphatidylcholine | AEA |
| 18:4(6Z, 9Z, 12Z, 15Z) phosphatidylethanolamine | AEA |
| 18:0/20:3(5Z, 8Z, 11Z) phosphatidylinositol-3-phosphate | 2-AG |

Furthermore, we have identified via Medscape significant up-regulation of 2-AG and the AEA metabolite ethanolamine (Figure 4), known for their analgesic properties^24^.

**Figure 4.**
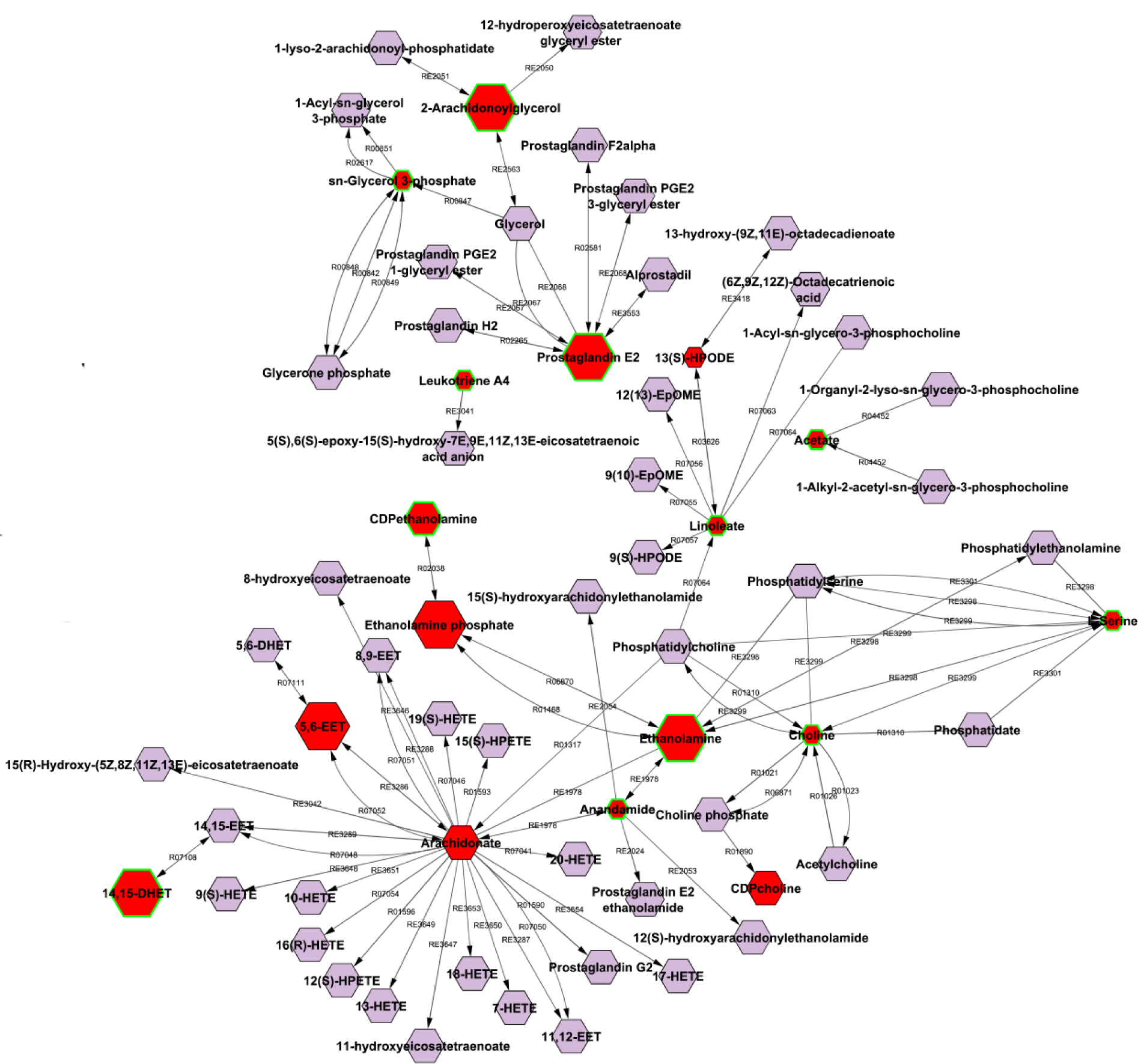
Infection with *L. mexicana* leads to enriched arachidonic acid metabolites. Normalized data from ear tissue of C57BL/6 mice infected with *L. mexicana* (12 weeks post infection) and naïve controls was used to perform integrative compound-compound network analysis with Metscape. Known anti-nociceptive metabolites from arachidonic acid metabolism, glycerophospholipid metabolism, linoleate metabolism, and prostaglandin formation from arachidonate are highlighted in the graph. Larger hexagons represent up-regulation, while smaller hexagons represent down-regulation. Red hexagons represent compounds detected in our data set, while hexagons with a green outline represent statistically significant metabolites.

Overall, our results show the presence of arachidonic acid and endocannabinoid metabolites with anti-nociceptive functions, which could contribute to the lack of pain experienced by CL patients.

### Immortalized and primary macrophages infected with *L. mexicana* display metabolic differences compared to naïve macrophages

Our initial screening revealed the upregulation of anti-nociceptive pathways at the infection site during *L. mexicana* infection. However, different cell types are present in the skin, and it is not yet known which one/s are responsible for secreting these metabolites. In order to determine a potential cell source, we performed untargeted mass spectrometry on infected macrophages, the canonical hosts for *Leishmania* parasites. First, we used RAW 264.7 macrophages, an immortalized cell line, and then validated our results with bone marrow-derived macrophages (BMDMs), in order to correlate our preliminary results with physiologically relevant primary cells (Fig. 5). We selected significant features, using volcano plots with a fold change threshold (x) 2 and t-tests threshold (y) 0.05 (Fig. 6 a and b), and identified unique molecular pathways enriched during *L. mexicana* infection, using partial least squares-discriminant analysis (PLS-DA) (Fig. 6 c and d). Not surprisingly, RAW 264.7 macrophages and BMDMs displayed distinct metabolic profiles in both the positive (Fig. 6 a and c) and negative (Fig. 6 b and d) modes, depending on their infection status.

**Figure 5.**
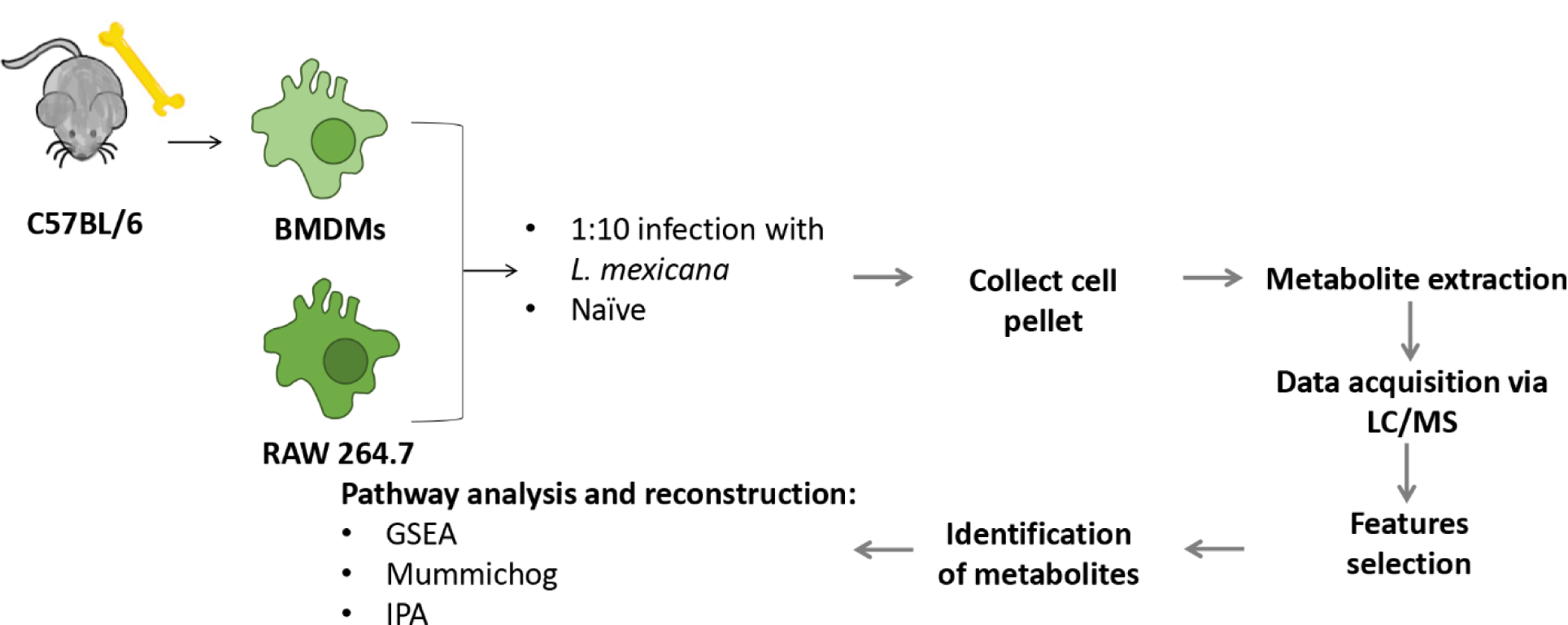
Experimental design for metabolomic analysis of analgesic pathways following *L. mexicana* infection. For bone marrow-derived macrophages (BMDMs), bone marrow was extracted from C57BL/6 mice and the cells were cultured in L-929 medium to allow for differentiation into macrophages. BMDMs and RAW 264.7 macrophages were the infected with stationary phase *L. mexicana* parasites or treated with media (naïve). After 24 hours, the cell pellets were collected and processed for mass spectrometry. The data acquired was used to identify enriched metabolites and determine activated pathways in the tissue. N=3 for each group.

**Figure 6.**
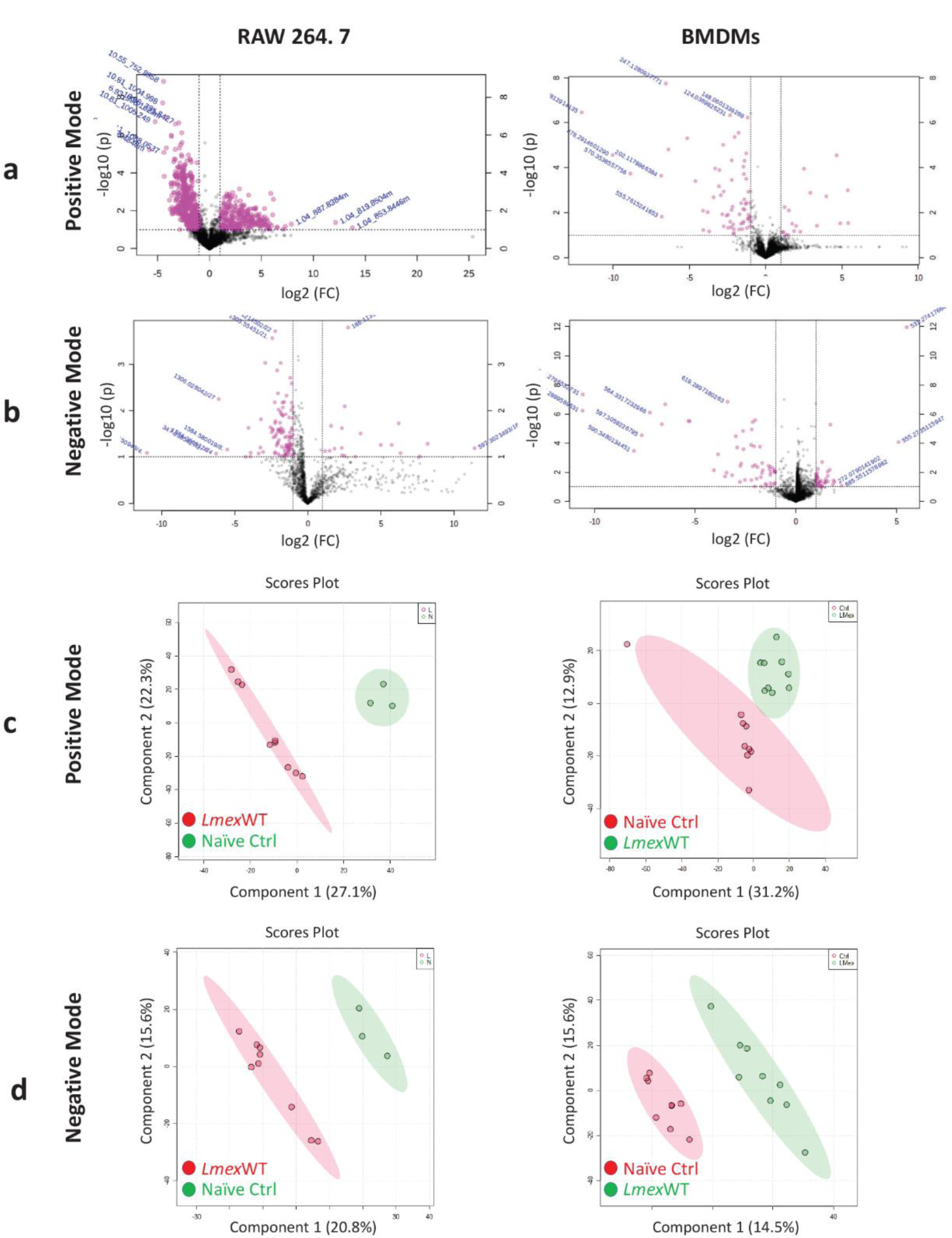
Infection with *L. mexicana* leads to unique metabolic signatures in macrophages. Normalized data from RAW 264.7 macrophages or C57BL/6-derived BMDMs infected with *L. mexicana,* and naïve controls was used to perform statistical analysis. a, b) Features selected by volcano plot from positive (a) and negative (b) modes for RAW 264.7 macrophages or C57BL/6-derived BMDMs cell pellets using LC/MS with log-transformed fold change threshold (2-fold, x-axis) and t-tests threshold (0.05, y-axis). c, d) PLS-DA from positive (c) and negative (d) modes for RAW 264.7 macrophages and C57BL/6-derived BMDM cell pellets.

### *L. mexicana* infection leads to enrichment of purine metabolism in immortalized and primary macrophages

We found enrichment of numerous metabolic pathways in infected RAW 264.7 macrophage and BMDMs, compared to uninfected controls, highlighting a dramatic metabolic shift during infection. In particular, we were interested in identifying any significantly enriched purine and arachidonic acid/endocannabinoid metabolites. Our results showed no arachidonic acid metabolism enrichment in RAW 264.7 macrophages and BMDMs at 24hrs post infection.

However, using the MetaboAnalyst software we found enrichment in caffeine metabolism in *L. mexicana*-infected RAW 264.7 macrophages (Fig. 7a). In particular, caffeine metabolism was the 6^th^ most enriched pathway in these cells (Fig. 7a). Further investigating this pathway with ingenuity pathway analysis (IPA), which uses compound IDs matched through different metabolic databases, we found that several purines analogous to caffeine metabolites were highly upregulated during infection. Specifically, we found a 17000-fold increase in nebularine, an adenosine analog not normally present at substantial levels in mammalian tissue, in infected compared to uninfected BMDMs (Fig. 7b). We also observed a 156-fold increase in xanthine, a 43-fold increase in inosine, and a 13-fold increase in hypoxanthine, which are all purines and derivatives of adenosine (Fig. 7b). Table 2 shows additional significantly upregulated purine metabolites in RAW 264.7 macrophages and BMDMs during *L. mexicana* infection based on mummichog and GSEA analysis. These results identify macrophages as a source of anti-nociceptive purine metabolites during infection, and potential players in *Leishmania*-mediated analgesia.

**Figure 7.**
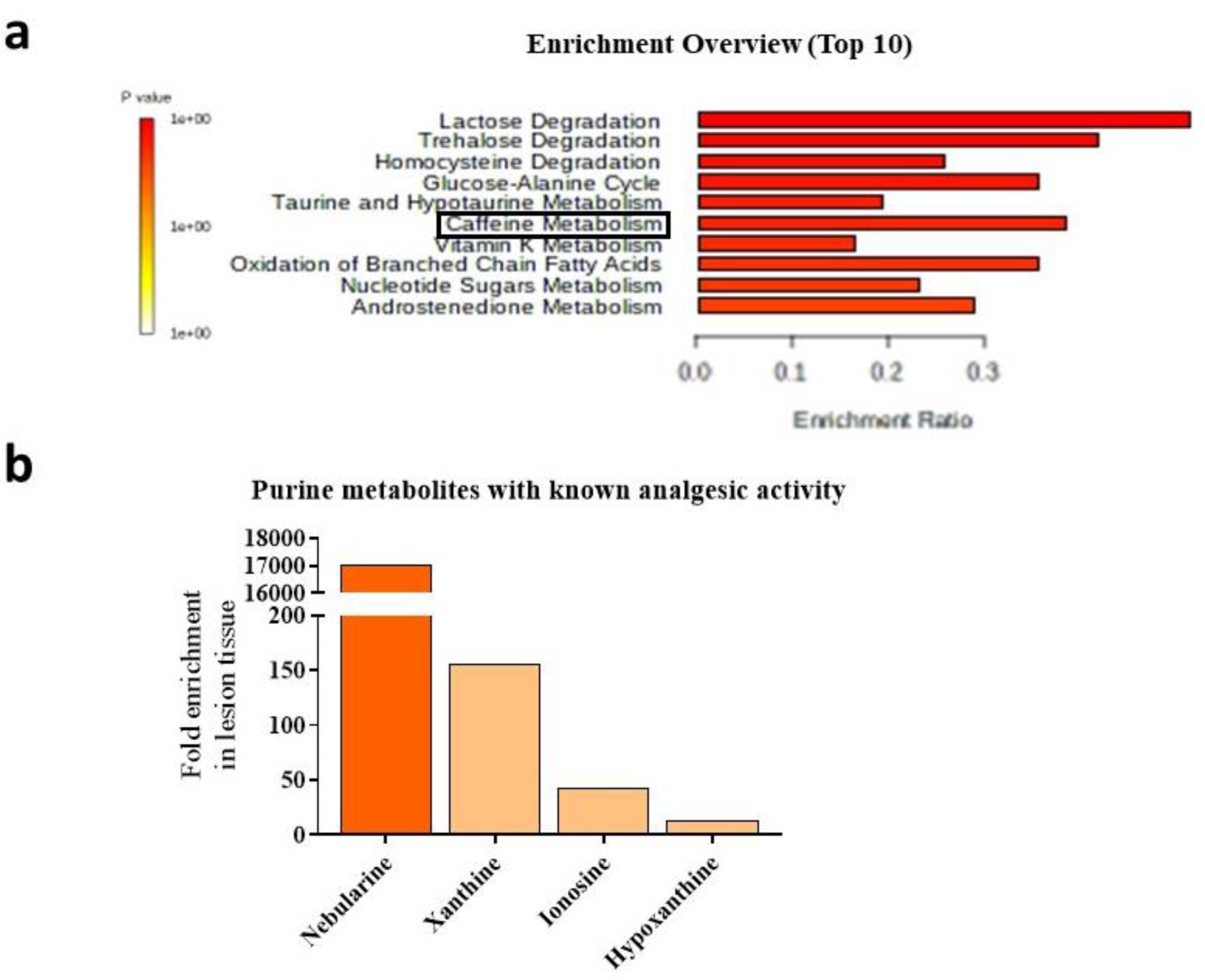
Infection with *L. mexicana* leads to purine metabolism enrichment in macrophages. Normalized data from RAW 264.7 macrophages **(a)** or C57BL/6-derived BMDMs **(b)** infected with *L. mexicana* and naïve control. **a)** Enrichment overview of top 10 enriched pathways in infected relative to naïve RAW cell pellets in the positive data set based on based on the MS Peaks to Paths module in MetaboAnalyst5.0. In the black box is highlighted the caffeine metabolism (6^th^ from the top). **b)** Fold change of most enriched caffeine metabolites in BMDMs infected with *L. mexicana* relative to naïve in the positive (dark orange) and negative (light tan) mode based on ingenuity pathway analysis (IPA).

**Table 2.** Significantly upregulated purine metabolites in macrophages during infection with **L. mexicana, compared naïve macrophages**. Normalized data from RAW 264.7 macrophages, and BMDMs infected with L. mexicana, or uninfected naïve controls was used to perform pathway analysis. Mummichog and GSEA analyses showed purine metabolites were significantly elevated in L. mexicana-infected RAW 264.7 macrophages and BMDMs compared to the respective uninfected naïve controls.

| RAW 264.7 macrophages | BMDMs |
| --- | --- |
| 1,7-Dimethylxanthine | 1,7-Dimethylxanthine |
| Theobromine; 3,7-Dimethylxanthine | 1-Methylxanthine |
|  | 1,3,7-trimethylurate; 8-oxy-caffeine |
|  | Theobromine; 3,7-Dimethylxanthine |
|  | 7-Methylxanthine |
|  | 3,6,8-Trimethylallantoin |
|  | 3,7-Dimethylurate |
|  | 1,7-Dimethylurate |
|  | 5-Acetylamino-6-formylamino-3-methyluracil |

### Anti- and pro-nociceptive mediators enriched in the *L. mexicana*-infected compared to naïve datasets

Other than caffeine/purine metabolism and arachidonic acid/endocannabinoid metabolism, our untargeted mass spectrometry analysis revealed up-regulation or down-regulation of several additional metabolites which are known to affect pain, based on previous studies. We have summarized in Tables 3 and 4 all the mediators with known anti-and pro-nociceptive properties respectively, statistically up-regulated or down-regulated in our *L. mexicana*-infected *in vitro* and *in vivo* data sets, compared to their naïve controls. For *in vivo* studies, we have also included an early infection model (7 days post infection).

**Table 3.**
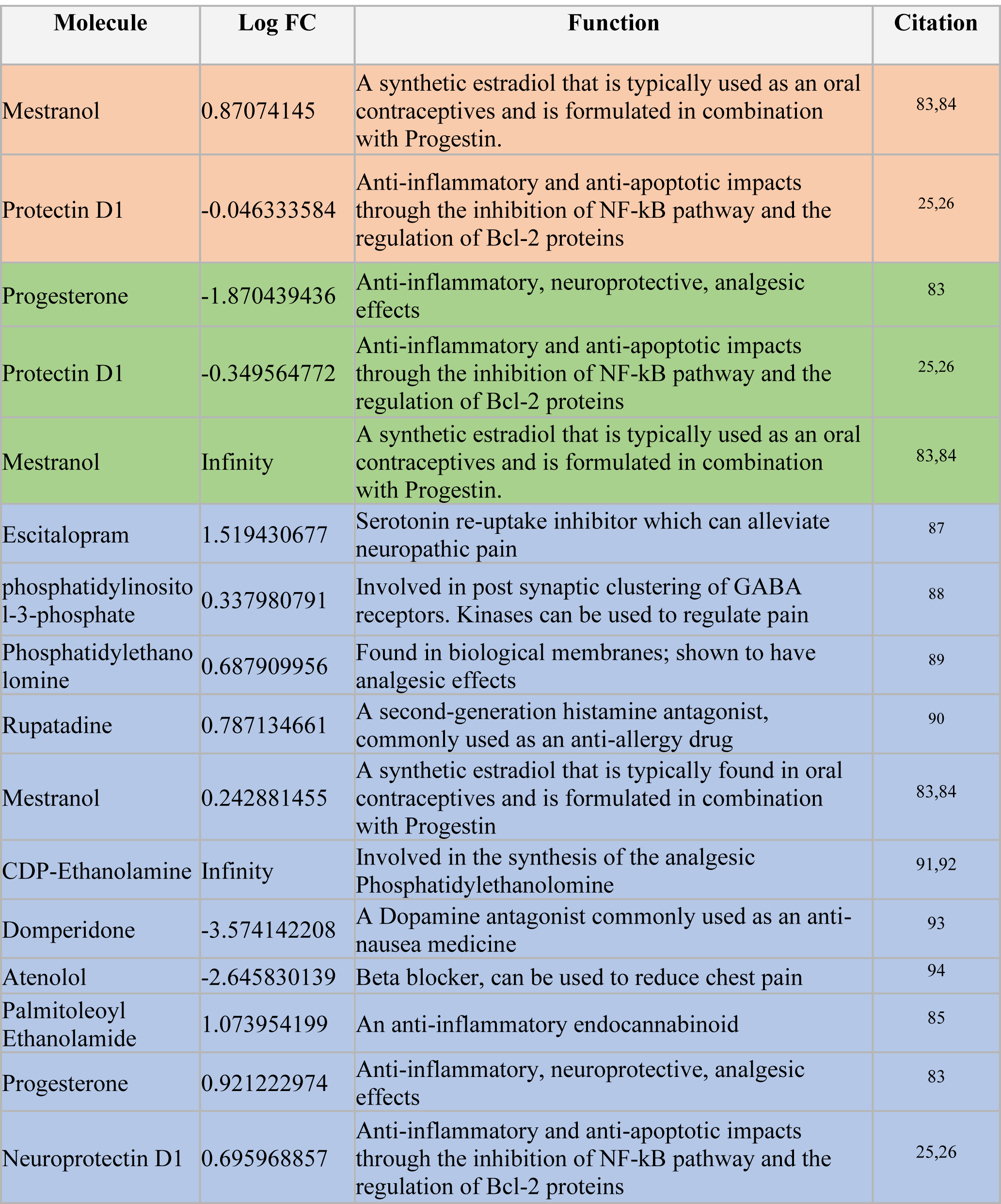
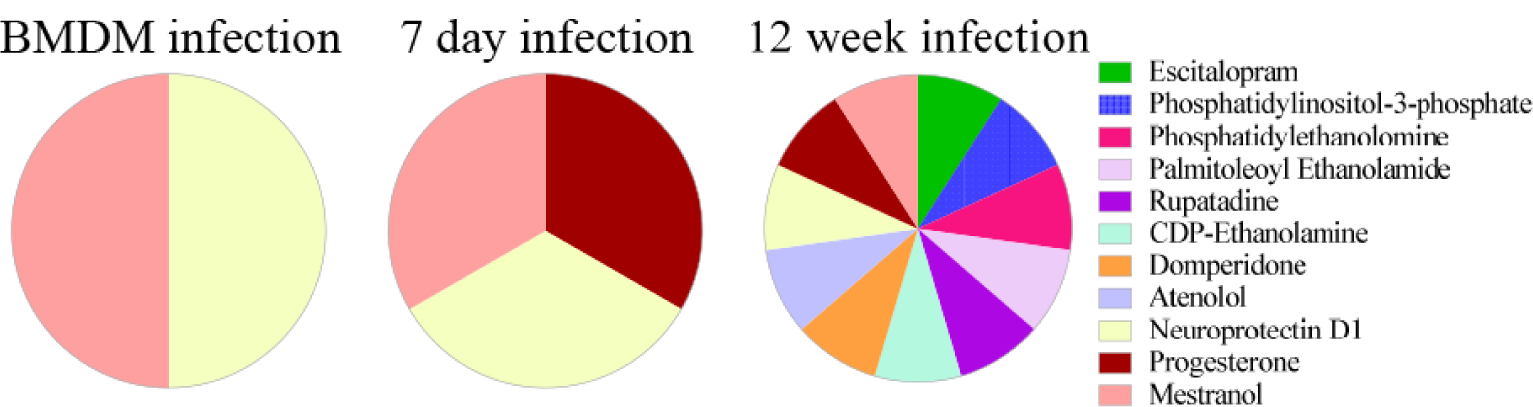
Anti-nociceptive mediators enriched in the *L. mexicana*-infection compared to naïve controls. Normalized data from BMDMs and ear tissue of C57BL/6 mice (7day and 12 week infection) infected with *L. mexicana* and naïve controls was used to identify anti-nociceptive metabolites reported in the literature. Log fold change represents enrichment of metabolites in the infected tissue/BMDM compared to uninfected controls. The table is color coded depending on the data set: orange represents metabolites found in the BMDM data set, green in the 7 day data set, and blue in the 12 week data set. The pie charts show anti-nociceptive metabolites shared between the three datasets.

**Table 4.**
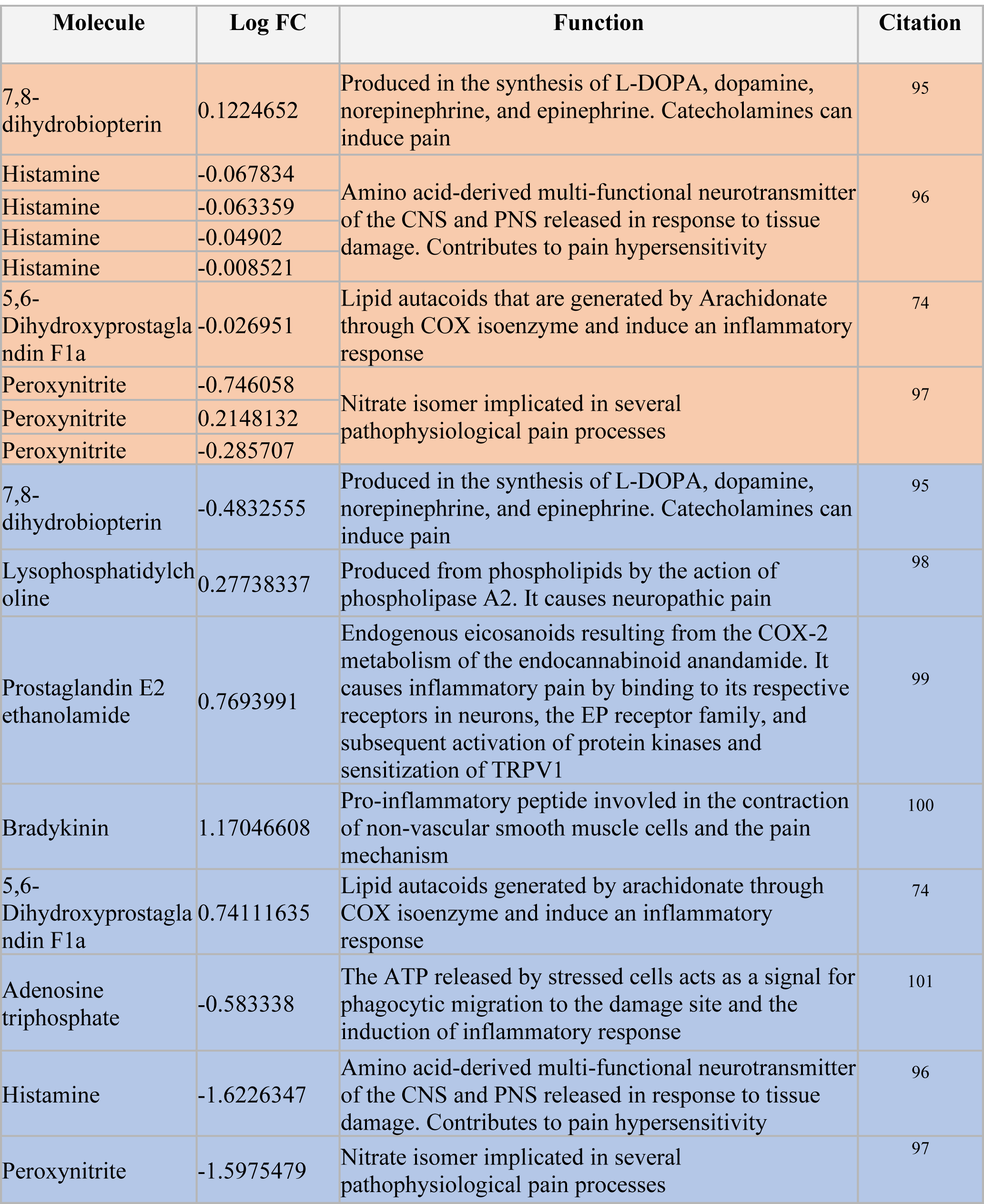
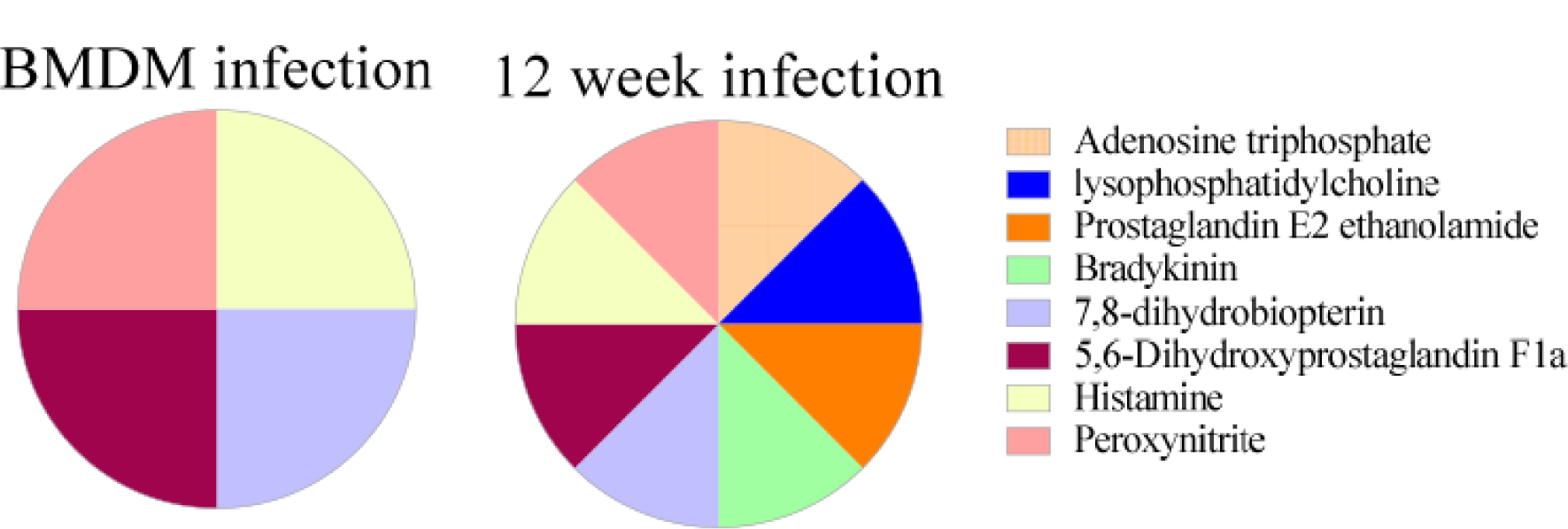
Pro-nociceptive mediators enriched in the L. mexicana-infection compared to naïve controls. Normalized data from BMDMs and ear tissue of C57BL/6 mice (12 week infection) infected with L. mexicana and naïve controls was used to identify pro-nociceptive metabolites reported in the literature. Log fold change represents enrichment of metabolites in the infected tissue/BMDM compared to uninfected controls. The table is color coded depending on the data set: orange represents metabolites found in the BMDM data set and blue in the 12 week data set. The pie charts show pro-nociceptive metabolites shared between the two datasets.

Overall, our results reveal the presence of metabolic mediators that can affect nociception at the lesion site and could play a role in analgesia experienced by CL patients.

## DISCUSSION

CL patients, including those infected by *L. mexicana*, routinely report analgesia in their cutaneous lesions^6–8^. Other pathogens are also known to induce painless skin lesions, such as *Mycobacterium ulcerans,* the causative agent of Buruli ulcer^36^, and *Treponema pallidum*, the causative agent of syphilis^37^. On the other hand, lesions caused by *Staphylococcus aureus*^9^, *Herpes* virus^10^, and *Varicella zoster* virus^11^ are painful. Taken together these observations indicate that pathogens may play a role in modulating nociception, either directly or indirectly, at the lesion site.

Several studies have been conducted in the context of *Leishmania* and other pathogenic infections to determine the mechanisms responsible for pain, or lack of, at the lesion site. In particular, previous literature has focused on the role of cytokines. Just as pain neurons, called nociceptors, express protein channels to directly detect painful stimuli, they also have receptors for cytokines, lipid mediators, and other molecules released by immune cells during both acute and chronic inflammation. The interaction of these mediators with receptors on nociceptors can modulate the sensitivity of the neuron making it more or less likely to fire an action potential and relay the pain signal to the spinal cord and finally the brain^22^. Extensive evidence show that pro-inflammatory cytokines such as IL-1β, IL-6, and TNF-α are involved in nociception^38^ and can be found in the painful *Staphylococcus aureus*^39^ and *Varicella zoster* virus^40^ infections. However, levels of IL-1β and TNF-α are also elevated in susceptible *L. mexicana* models^41, 42^, usually associated with lack of pain at the lesion site. This contradiction is accompanied by conflicting results found by previous studies in pre-clinical models of *L. major*-mediated anti-nociception^16–21^. The cytokine milieu, as well as the pain response in these studies varied greatly depending on the mouse model, inoculum, time point, and other factors^16–21^. Taken together these observations suggest that cytokines do not provide a complete picture of which mechanisms are responsible for CL-mediated analgesia, and that other mediators might be involved.

After inoculation, *Leishmania* parasites reside within phagocytes such as macrophages^1^. It is well known that *Leishmania* can alter the metabolic profiles of infected cells^43–45^ and this can modulate immunological functions in the host to promote parasitic survival^43, 45–47^. However, the impact of this metabolic reprogramming on the lack of pain experienced by CL patients at the lesion site is unclear. In this study we were interested in using metabolomics approaches to investigate whether metabolic changes during the *L. mexicana* infection can modulate pain in cutaneous lesions. To our knowledge, the current study is the first to use metabolomics to investigate the relationship between metabolic and immunological changes in the context of pain during leishmaniasis. We found the enrichment of several pathways with known anti-nociceptive properties, in mice chronically infected with *L. mexicana,* as well as in infected macrophages *in vitro*.

In particular, we found that several endogenous purines, which are analogous to metabolites produced by the caffeine metabolism, were highly upregulated during infection. Before proceeding in our investigation of CL-mediated changes in purine metabolism, we confirmed with our veterinary protocols that there was no caffeine enrichment of the mouse chow. *Leishmania* parasites are purine auxotrophs and scavenge purine from the host cell in order to replicate^48, 49^, therefore it is not surprising that infected host cells upregulate synthesis of purine metabolites such as the ones identified during infection *in vitro* and *in vivo*. In particular,

*L. mexicana* parasites are known to utilize purine metabolites such as xanthine, hypoxanthine, and ionosine as an energy source for cellular functions and replication^50^. Purine metabolites released by *L. mexicana*-infected macrophages could have modulatory paracrine functions on neighboring cells at the lesion site, as these metabolites are known to have multiple anti-inflammatory and immunomodulatory properties^51–55^. For instance, purines can induce an M2 immunosuppressive phenotype in macrophages, characterized by production of the cytokine IL-10, which is known to have anti-inflammatory and analgesic properties^56, 57^. The role of IL-10 in susceptibility to *L. mexicana* has been previously described^58, 59^, and this cytokine has been associated with hypoalgesia in *L. major*-infected BALB/c mice^16^. Furthermore, it is possible that in addition to promoting an anti-inflammatory environment permissive to *Leishmania* survival and replication, these purines could directly modulate nociception at the lesion site due to their action on adenosine receptors (ARs) on nociceptive neurons. The role of nebularine, traditionally identified as a fungal metabolite, and hypoxanthine in pain is unknown. However, xanthine derivatives^55^ and ionosine^54^ have been shown to have anti-nociceptive properties by acting through the ARs. ARs are key regulators of inflammation and pain in nociceptors and other cells. In particular, signaling through A1R and A3R leads to a decrease in cAMP levels^51^ and contributes to analgesia^60–63^. Purine metabolites can act on A1R and A3R to decrease nociception in sensory neurons by inhibiting excitatory Ca^2+^ currents^60–62^, and to impair transient receptor potential vanilloid (TRPV)-1 channel function indirectly by depleting PiP2 via A1R signaling, or directly by binding to TRPV1 channels^60, 61, 63^. TRPVs are non-selective Ca^2+^ channels responsible for sensing a wide range of painful stimuli and can mediate anti-nociception when inactivated or downregulated^64, 65^. Taken together, these observations suggest that upregulated purine metabolism can result in increased production of anti-nociceptive cytokines, such as IL-10, and can also directly modulate pain at the lesion site by interacting with nociceptive neuronal receptors. However, these purine metabolites might not be the only ones involved in CL-mediated analgesia.

Further analysis of upregulated mediators with known analgesic properties revealed upregulated metabolites of the arachidonic acid pathway at the lesion site of infected mice, compared to naïve mice. Arachidonic acid is a metabolite of anandamide (AEA) and 2-arachidonoylglycerol (2-AG), the two known endocannabinoids^66^. These functional lipids act on the cannabinoid receptors (CB) 1-2 to modulate different functions including inflammation and pain^24^. CB1s reside throughout the central nervous system, while CB2s are mainly found in the peripheral nervous system and on immune cells^67^. Action through CBs leads to anti-nociception and suppression of pro-inflammatory cytokine production^68^. AEA specifically is a ligand for both CB1/CB2 and TRPV1, causing desensitization of TRPV1 at high concentrations^69^. Topical administration of cannabinoid compounds has been used for a long time to treat pain and has been validated in pre-clinical and clinical studies^70–72^. Furthermore, there is already evidence of *Leishmania* infection modulating arachidonic acid metabolism. In particular, infection with *L. donovani*, a visceral leishmaniasis strain, results in increased levels of arachidonic acid in murine peritoneal macrophages^73^. Arachidonic acid is also the precursor of other biosynthetic pathways, such as the production of prostaglandins, thromboxane, and lipoxins which have been implicated in modulating inflammation and pain^74, 75^. While prostaglandins and thromboxane can promote inflammation and nociception^74, 76^, lipoxin A4 can lead to anti-nociception^75^.

Based on these observations we hypothesize that *L. mexicana* infection can reduce inflammation and pain at the lesion site by upregulating the production of arachidonic acid and endocannabinoid metabolites, resulting in the development of a parasite-permissive environment. The anti-nociceptive properties of endocannabinoids have been implicated not only in inflammatory pain, but also in chronic pain^77^, which could be a beneficial mechanism for *Leishmania* to utilize for long term survival.

Interestingly, while we have shown upregulation of purine metabolism *in vitro*, we did not observe enriched arachidonic acid metabolism in macrophages infected with *L. mexicana*. These results suggest that macrophages might not be the major source of arachidonic acid/endocannabinoid metabolites in CL lesions, or that the *in vitro* nutrient milieu may not fully reflect the *in vivo* conditions. Arachidonic acid is found in the cell membrane phospholipids of many different cell types and stored in the lipid bodies of endothelial cells, fibroblasts, and immune cells such as macrophages, neutrophils, mast cells, and eosinophils^78, 79^.

Endocannabinoids are produced by neurons, as well as immune cells such as macrophages, basophils, and activated T and B cells^80–82^. Based on these observations, further research is necessary to determine the main source of arachidonic acid and endocannabinoid mediators in CL lesions. Additionally, more studies are needed to determine whether the analgesic metabolites identified in this study are also present in physiologically relevant levels in human *L. mexicana* lesions, and whether they significantly contribute to the lack of pain experienced by CL patients.

In this study we have focused on purine and arachidonic acid metabolism due to their already established analgesic properties. However, our metabolomics analysis also revealed the enrichment of other pathways such as steroid hormones biosynthesis, catecholamine biosynthesis, nicotinate metabolism, and sphingolipid biosynthesis, which have also been implicated in pain modulation to some degree. In particular, mestranol was upregulated in infected samples across all data sets. Mestranol is a synthetic estrogen analog and has been implicated in anti-nociception^83, 84^. While mestranol was not administered in our experiments, it is possible that an analogous estrogen was identified as mestranol by MetaboAnalyst. Mice chronically infected with *L. mexicana* also showed upregulated palmitoleoyl ethanolamide, an endocannabinoid^85^; progesterone, an endogenous steroid^83^; and protectin D1^25, 26^, a pro-resolving lipid with an anti-inflammatory and anti-nociceptive properties. Interestingly, some of the anti-nociceptive mediators were also downregulated in infected samples, as shown by a negative LogFC. Furthermore, our analysis revealed the presence of pro-nociceptive mediators in our infected data sets, several of which, including histamine, were downregulated in infected samples. Taken all together, these results show a balance between pro-and anti-nociceptive mediators at the lesion site. Our future studies aim to elucidate the contribution of these pathways to analgesia in CL and determine whether they can interact synergistically to create an anti-inflammatory and anti-nociceptive environment at the lesion site.

Overall, this study provides the first evidence of metabolic pathways upregulated in *L. mexicana* infection that have a known molecular basis for anti-nociceptive effects experienced by individuals infected with the parasite.

## MATERIALS AND METHODS

### Mouse strains and parasites

Female C57BL/6 mice were purchased from Envigo (Harlan laboratories) Indianapolis, IN, USA, and housed at The Ohio State University animal facility, following approved animal protocols and University Laboratory Animal Resources (ULAR) regulations (2010A0048-R3 Protocol). All the experiments were performed using 5 age-matched 5-8 week old female mice per group.

129S6/SvEvTac mice (purchased from Taconic Biosciences, Inc.) were used to maintain *L. mexicana* (MNYC/B2/62/m379) parasites via subcutaneous inoculation into the shaved back rumps. Amastigotes were then obtained from the draining lymph nodes of infected mice and grown at 26°C in M199 medium supplemented with 1% Penicillin/Streptomycin, and 1% HEPES10% fetal bovine serum (FBS), to generate stationary phase promastigotes.

### *In vitro* cell culture and infection

For *in vitro* studies we used the immortalized RAW 264.7 macrophage cell line, as well as primary bone marrow-derived macrophages (BMDMs). The RAW 264.7 cell line was purchased from American Type Culture Collection (ATCC), and cell identity was verified regularly based on cell morphology. Both RAW 264.7 and BMDMs were cultured with RPMI medium supplemented with 10% fetal bovine serum (FBS), 1% penicillin/streptomycin and 1% HEPES at 37°C with 5% CO2. BMDMs were obtained from the femur and tibias of C57BL/6 mice. After isolation, the bone marrow was cultured with complete RPMI supplemented with 20% supernatant from L-929 cells for 7-10 days until differentiation was complete. RAW 264.7 macrophages and BMDMs were plated in a 24-well plate at a density of 0.5x10^6^ per well and infected overnight with stationary phase *L. mexicana* promastigotes at a ratio of 10:1 (parasite to macrophages). The controls were treated with media alone. Then, the extracellular parasites were removed by washing with PBS and new media was applied. After a 24hrs incubation, the supernatant and cell pellet were collected and processed for mass spectrometry.

### *In vivo* infection

Aged-matched C57BL/6 mice were inoculated intradermally in the ear with 1 × 10^6^ *L. mexicana* promastigotes in the stationary phase. After 7 days or 12 weeks, the ipsilateral infected ear, and the naïve ears were collected and processed for mass spectrometry. For RT-PCR analysis, the ear samples were collected at 3, 6, 9, and 12 weeks post infection and placed in RNAlater (Thermo Fisher Scientific, Waltham, MA) until further processing.

### Mass spectrometry

For *in vitro* experiments, the culture supernatant was collected and cell debris was removed by centrifugation according to SOP 5 of the Laboratory Guide for Metabolomics Experiments ^86^. The attached cells were be scraped, washed with PBS and quenched with 80% methanol. Then they were snap-frozen, centrifuged, and lyophilized according to SOP 4 of the Laboratory Guide for Metabolomics Experiments ^86^.

For *in vivo* studies, the ears were collected, snap frozen, and processed for mass spectrometry analysis according to SOP 7 of the Laboratory Guide for Metabolomics Experiments ^86^. Samples were then incubated with 500 uL of 100% MeOH and sonicated. The tissue was weighed and homogenized at 40 mg/mL of 50% MeOH solution for 3 cycles in a homogenizer with Precellys. The supernatant was collected, dried down, and reconstituted in ½ of the original volume in 5 % MeOH with 0.1 % formic acid.

Untargeted analysis was performed on a Thermo Orbitrap LTQ XL with HPLC separation on a Poroshell 120 SB-C18 (2 x 100 mm, 2.7 µm particle size) with an WPS 3000 LC system. The gradient consisted of solvent A, H2O with 0.1 % Formic acid, and solvent B 100 % acetonitrile at a 200 µL/min flow rate with an initial 2 % solvent B with a linear ramp to 95 % B at 15 min, holding at 95% B for 1 minutes, and back to 2 % B from 16 min and equilibration of 2 % B until min 32. For each sample, 5 µL were injected and the top 5 ions were selected for data dependent analysis with a 15 second exclusion window.

For feature selection in the untargeted results analysis, including database comparison and statistical processing, samples were analyzed in Progenesis QI and the pooled sample runs were selected for feature alignment. Anova p-value scores between the groups were calculated with a cutoff of < 0.05. With database matching using the [Human Metabolome Database], selecting for adducts M+H, M+Na, M+K, and M+2H and less than 10 ppm mass error, unique features were tentatively identified as potential metabolites.

### Statistical analysis of mass spectrometry datasets

Peak intensity data tables from the mass spectrometry experiment were formatted into comma-separated values (CSV) files conforming to MetaboAnalyst’s requirements and uploaded into the one-factor statistical analysis module. Each analysis passed MetaboAnalyst’s internal data integrity check and additional data filtering was performed based on interquartile range. On the normalization overview page, sample normalization was performed based on the median of the data and the auto-scaling option was chosen to perform data scaling; No transformation of the data was performed. For dimensionality reduction, both principal component analysis (PCA) and partial least-squares discriminant analysis (PLSDA) were employed. Cross-validated sum of squares (Q2) performance measures were used to determine if PLSDA models were overfitted.

Visualization of significant, differentially regulated metabolites was done by generating volcano plots with cutoffs of < 0.05 false-discovery rate (FDR) and > 2-fold change (FC). Clustering of samples and features were analyzed by creating dendrograms and hierarchical heatmaps, respectively.

### Pathway analysis of mass spectrometry datasets

We have used two different techniques in order to identify enriched pathways in our data sets. First we used the Functional Analysis Module (MS peaks to pathways) in MetaboAnalyst 4.0. Detected peaks (mass-to-charge ratios + retention times) from positive and negative analytical modes of the mass spectrometer for each sample were organized into four column lists along with calculated p-values and t-scores from univariate t-tests. These peak list profiles were uploaded to the functional analysis module and passed the internal data integrity checks. The ion mode in MetaboAnalyst was set to the appropriate type depending on the analytical mode that was used to generate the data. For each analysis the mass tolerance was set to 10 ppm, the retention time units were set to minutes, and the option to enforce primary ions was checked. In parameter settings, the mummichog algorithm (version 2.0) and the modified gene set enrichment algorithm were used for all analyses. The p-value cutoff for the mummichog algorithm was left at the default (top 10% of peaks). Currency metabolites and adducts were left at default settings. Lastly, the Kyoto Encyclopedia of Genes and Genomes (KEGG) pathway library for *Mus musculus* was selected as the metabolic network that the functional analysis module would use to infer pathway activity and predict metabolite identity; only pathways/metabolite sets with at least three entries were allowed.

To confirm the enriched pathways identified with MetaboAnalyst, we also used the Ingenuity Pathway Analysis (IPA) software. Metabolite matches to the detected peaks from database searches (HMDB and LIPID MAPS), along with calculated p-values and fold changes were uploaded into IPA software for core analysis. The reference set was selected as the Ingenuity Knowledge Base (endogenous chemicals only) and direct and indirect relationships were considered during analysis. Settings in the networks, node type, data sources, confidence, and mutations tabs were left at default values. The species tab settings were set to mouse and uncategorized (selecting uncategorized species for metabolomics is necessary in IPA as most metabolites are not unique to any one species). Lastly, in the tissues and cell lines tab all tissues were considered in the analysis, whereas all cell lines were excluded from consideration. The p-value cutoff for every analysis was set as 0.05 and the fold change cutoffs were adjusted to obtain between 200-1000 analysis-ready molecules and then kept the same across different analyses.

### Integrative Network Analysis

The Metscape 3.1.3 App from the Cytoscape 3.9.1 software was used in order to build integrative network analysis of our in-vivo dataset using Metscape’s internal database which incorporates KEGG and EHMN data. The IDs of the metabolomic dataset were converted from HMDB IDs to KEGG IDs recognized by the Cytoscape software via the Chemical Translation Service (CTS) and verified by the Metaboanalyst Compound ID Conversion tool. The metabolomic dataset was then uploaded as a compound file to Metscape and the P. Value and FC Ratio cutoff points were set at 0.05 and 1.0 respectively. 810 from a total of 2771 metabolites were not accepted as input compounds by Metscape and were removed from the dataset.

A Compound-compound network was created for the detected Purine Metabolism pathway of our dataset, along with an integrative network build from the combination of Arachidonic Acid Metabolism, Glycerophospholipid metabolism, Linoleate metabolism, and Prostaglandin formation from arachidonate. The two created networks allowed us to visualize the integrated relationship among the metabolites and Endocannabinoids involved in Purine metabolism and Arachidonic Acid metabolism respectively.

### Statistical analysis

All *in vitro* and *in vivo* data show a representative experiment with N ≥ 3 per group. N represents different biological replicates. For the mass spectrometry data all statistical analysis were performed with MetaboAnalyst and IPA softwares.

## DATA AVAILABILITY STATEMENT

All relevant data is available in the main text and supplementary information. Any additional information can be provided upon reasonable request to the authors.

## AUTHOR CONTRIBUTIONS

G.V., S.G., H.L.N., and A.R.S. designed the experiments. G.V., B.C., and Y.M. performed the experiments. T.O. and N.A. analyzed the mass spectrometry data. G.V. and S.G. wrote the manuscript. G.V., T.O., B.C., Y.M., N.A., C.A., S.G., H.L.N, and A.R.S. revised the manuscript.

## Abbreviations

AR: adenosine receptor
TRPV1: transient receptor potential channels of the vanilloid subtype 1
IL-10: interleukin-10
PiP2: Phosphatidylinositol 4,5-bisphosphate
CB: cannabinoid receptor

